# Tuning network dynamics from criticality to the chaotic balanced state

**DOI:** 10.1101/551457

**Authors:** Jingwen Li, Woodrow L. Shew

**Affiliations:** Department of Physics, University of Arkansas, Fayetteville, Arkansas, United States of America

## Abstract

According to many experimental observations, neurons in cerebral cortex tend to operate in an asynchronous regime, firing independently of each other. In contrast, many other experimental observations reveal cortical population firing dynamics that are relatively coordinated and occasionally synchronous. These discrepant observations have naturally led to a lively debate surrounding discrepant hypotheses. A commonly hypothesized explanation of asynchronous firing is that excitatory and inhibitory neurons are precisely correlated, nearly canceling each other, resulting in the so-called ‘chaotic balanced state’. On the other hand, the ‘criticality’ hypothesis posits an explanation of the more coordinated state that also requires a certain balance of excitatory and inhibitory interactions. Both hypotheses claim the same qualitative mechanism - properly balanced excitation and inhibition. Thus, a natural question arises: how are the chaotic balanced state and criticality related, how do they differ? Here we propose an answer to this question based on investigation of a simple, network-level computational model. We show that the strength of inhibitory synapses relative to excitatory synapses can be tuned from weak to strong to generate a family of models that spans a continuum from ‘criticality’ to the ‘chaotic balanced state’. Our results bridge two long-standing competing hypotheses and offer a possible explanation of discrepant experimental observations: neuromodulatory mechanisms that tune the strength of excitatory and inhibitory synapses may tune cortical state from criticality to the balanced state, and to intermediate states between these extremes.

**Author summary:** What is the dynamical state of cerebral cortex? Are neurons mostly uncorrelated, firing asynchronously with each other? Are synchronous oscillations important? The answers to these questions have fundamental implications for how the cortical neural population encodes and processes information. Here we show that two possible scenarios - criticality and the chaotic balanced state - that are typically considered incompatible can be attained in the same network by maintaining a certain kind of balance, while tuning the strength of inhibition relative to excitation.

## Introduction

Mounting experimental evidence supports the hypothesis that the cerebral cortex operates in a dynamical regime near criticality [1–8]. What do we mean by criticality? Often, criticality is described as a boundary in the space of possible dynamical regimes. On one side of the boundary population activity tends to be orderly, correlated across the entire network. On the other side, neurons fire more independently of each other. At criticality, population dynamics are more diverse, rarely exhibiting synchronization that spans the network, but often showing coordinated firing among groups of neurons at small and intermediate scales [9, 10]. Direct evidence that the cerebral cortex may indeed operate near such a boundary comes from experiments and models in which the balance of excitation (E) and inhibition (I) is disrupted. These studies show that one can push cortical dynamics from a dynamical regime consistent with criticality to a hyperactive synchronous regime by suppressing inhibiton (GABA antagonists) or to a low-firing asynchronous state by increasing inhibition (GABA agonists) [11–15]. Similarly, critical dynamics can be pushed to a low-firing asynchronous regime by suppressing excitation (AMPA and NMDA antagonists) [13–15]. These observations support the hypothesis that the cortex may operate near criticality under normal conditions, but only if the proper balance of E and I is maintained.

However, not all observations of the cortex under ‘normal conditions’ exhibit the diverse multi-scale coordination that is expected near criticality. Indeed, many experimental measurements have revealed relatively asynchronous firing, particularly in vigilant and active behavioral conditions [16–20]. One of the most prominent theoretical explanations of the asynchronous state, the so-called ‘chaotic balanced state’ hypothesis, is based on balanced E and I [21–24]. The idea is that E and I inputs to any given neuron wax and wane together, nearly canceling each other most of the time. Some experiments support this possibility based on whole cell recordings of E and I inputs [25–27]. During brief moments the E-I cancellation is imperfect and neurons can fire.

Both the criticality hypothesis and the chaotic balanced state hypothesis seem to require balanced E and I. Support for both hypotheses has been shown in awake animals. However, the difference in coordination of population activity for criticality versus the chaotic balanced state is stark. How can we reconcile these two hypotheses? When should we expect to see the coordination of criticality; when should we expect to see the asychronous activity of the chaotic balanced state?

Here we hypothesize that criticality requires a different kind of E/I balance than the chaotic balanced state. Here we address this possibility using a network-level model of probabilistic, binary neurons. By tuning a single parameter, we found that we can generate a family of models, spanning a continuum from criticality to the chaotic balanced state. When synapses are strong and balanced, the chaotic balanced state results. When synapses are relatively weak and balanced, criticality results. Our results offer a possible explanation for the variety of experimental observations, suggesting that the cortex could shift its dynamical regime from near criticality to the chaotic balanced state and a continuum of intermediate states between these extremes, all while maintaining balanced excitation and inhibition.

## Results

We study a recurrent network of *N* = 1000 probabilistic integrate-and-fire of binary neurons. There are 800 excitatory neurons and 200 inhibitory neurons. By altering excitatory and inhibitory interactions, we consider a range of different dynamical regimes. We tune the relative strengths of excitatory and inhibitory synapses including a range of I/E weight ratios between 0 and 4.6. For every I/E ratio, the synaptic weight matrix is normalized such that its largest eigenvalue is unity in magnitude. This constraint ensures that activity does not tend to grow nor decay on average as time passes [28, 29]. In this sense, the activity is stable for all the dynamical regimes we consider, but the stability is achieved by different combinations of E and I. In some previous work on criticality, the constraint of unity largest eigenvalue has been used to define ‘balance’ of excitation and inhibiton, but this should not be confused with the different meaning of ‘balance’ considered in previous studies of the chaotic balanced state.

Typical population dynamics of the system under different I/E ratios are shown in Fig 1a. When the I/E weight ratio is low, population activity is rather synchronized. The population tends to fire in bursts of coordinated activity with diverse burst sizes, separated by periods of quiescence. As the I/E weight ratio increases to moderate levels (I/E≈2), the network becomes less synchronized, and counter-intuitively, the population firing rate increases as the inhibition becomes stronger. However, this is consistent with previous work showing that inhibitory neurons in a recurrent network can make the dynamics ceaseless, provided that the largest eigenvalue of the weight matrix is unity [30]. As we increase the I/E weight ratio further, the network is further desynchronized and the population firing rate eventually decreases. As I/E is tuned from 0 to 4.6, synapse strength increases due to imposing the constraint that the largest eigenvalue of the synaptic weight matrix is unity (Fig 1b). Thus, the synchronized regime is characterized by weak, balanced synapses, while the asynchronous regime is characterized by strong, balanced synapses.

**Fig 1.**
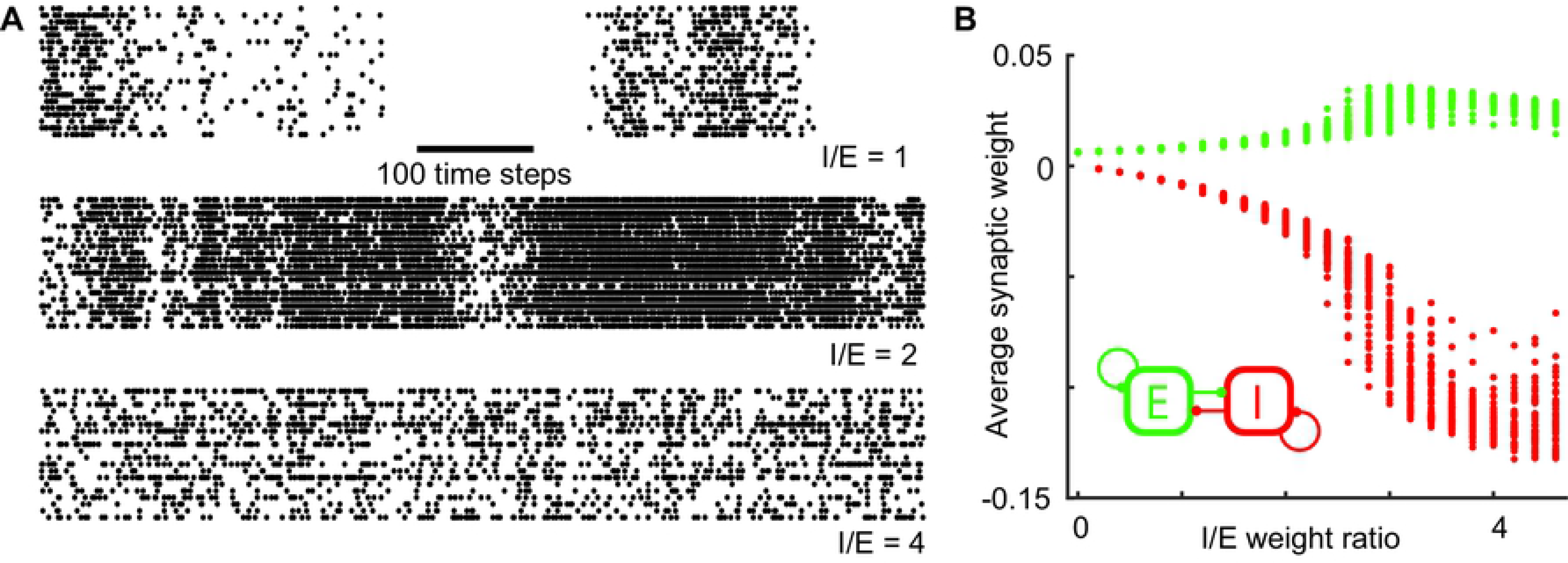
Balanced networks with stronger inhibition relative to excitation are more asynchronous. **(a)** Example spike raster plots are shown for I/E weight ratios of 1 (top), 2 (middle) and 4 (bottom). Population-level coordination decreases dramatically, reaching an asynchronous regime as inhibition is strengthened relative to excitation. **(b)** Average excitatory (green) and inhibitory (red) synaptic weight as a function of the I/E weight ratio for each realization. Synapse strength increases as I/E ratio increases due to the constraint of unity largest eigenvalue.

To more carefully examine the dynamics of excitation and inhibition, we consider separately the excitatory and inhibitory inputs to the model cells. For all I/E ratios, E and I are correlated, but as we increase the I/E ratio, the dynamics are tuned from a state in which excitatory input dominates (is not canceled by inhibition) to a state in which inhibitory input cancels the excitatory input more and more exactly (Fig 2a). We define ‘E/I tension’ to measure how tightly the excitatory and inhibitory inputs balance each other (Methods). Fig 2b shows that the E/I tension gradually increases as the I/E weight ratio increases, and reaches as high as 1 when the I/E weight ratio is 4. This result is consistent with previous work showing that tightly balanced excitation and inhibition that cancel each other leads to desychronization [23].

**Fig 2.**
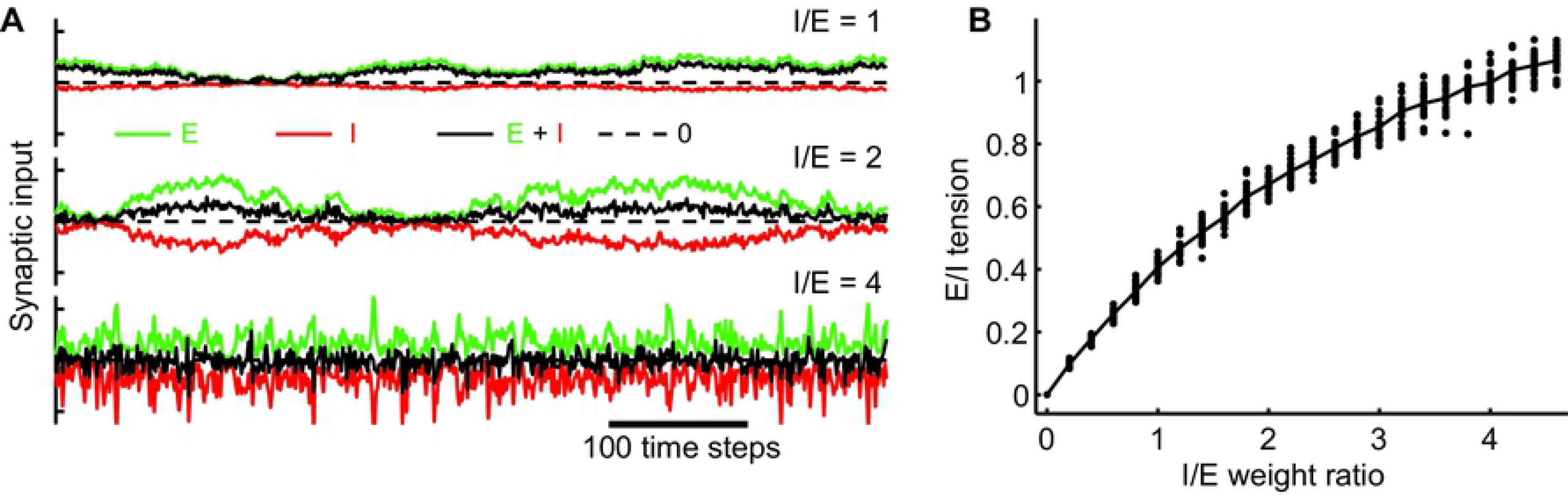
Excitatory and inhibitory inputs are more tightly balanced for larger I/E. **(a)** Synaptic inputs when the I/E weight ratios are 1 (top), 2 (middle) and 4 (bottom). The excitatory (green) and inhibitory (red) inputs are shown seperately from the total synaptic input (black). **(b)** E/I tension as a function of the I/E weight ratio. Dots are the values of each realization and the solid line is the mean value.

One way to test the criticality hypothesis is to examine distributions of neural avalanche sizes [11]. Similar to previous work [11, 30, 31], we define an avalanche as a period of time during which the number of active neurons exceed a threshold (Fig 3a). The duration and size of an avalanche are defined as the number of time steps and the total number of spikes that occurred during the avalanche, respectively. We found that both avalanche duration and size distributions were sensitive to changing I/E (Fig 3b). When the I/E weight ratio is low, the avalanche duration and size distributions are close to power-law distributions, as expected at criticality. As the I/E weight ratio increases, the distributions of avalanche duration and size deviate from power-law distributions, with large avalanches becoming less prominent. We use a previously developed measure, called *κ_ϵ_*, to quantify how much a distribution deviates from a power-law distribution with exponent *ϵ* [5, 11, 14]. If the distribution is close to the power-law distribution, then *κ*_*ϵ*_ is close to 1, which occurs for both avalanche duration and size distributions when the I/E weight ratio is low. Any deviation in *κ*_*ϵ*_ from 1 means a deviation from the power-law distribution. As the I/E weight ratio grows larger, *κ*_*ϵ*_ starts to deviate from 1, then varies erratically for intermediate I/E before settling near *κ*_*ϵ*_ = 0.8 as I/E approaches 4. The avalanche distributions indicate that the recurrent network exhibits dynamics consistent with criticality only when I/E weight ratios are low, i.e., weak inhibition.

**Fig 3.**
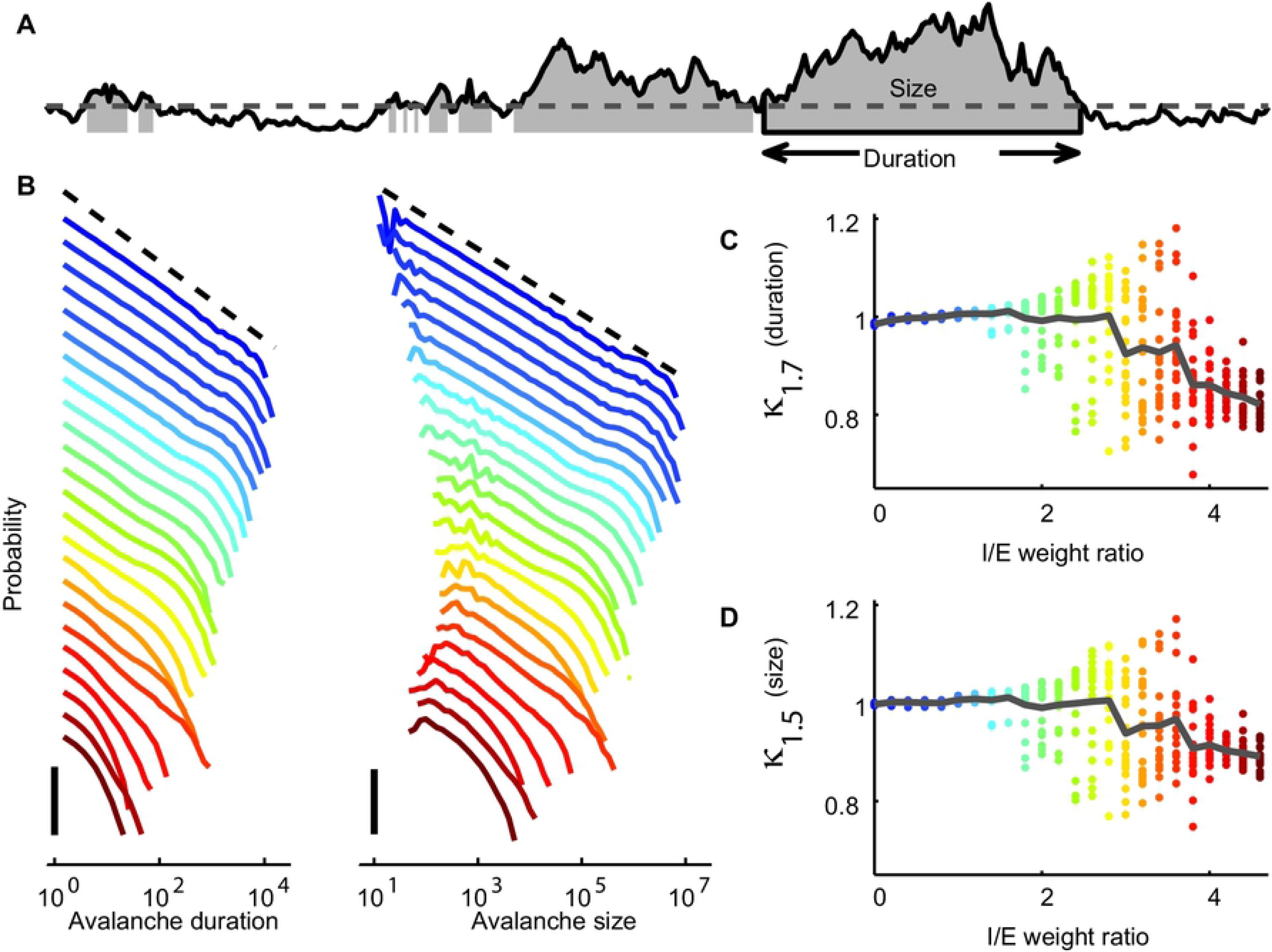
Avalanche distributions consistent with criticality only when the I/E weight ratio is low. **(a)** An avalanche is defined as a time period where the number of active neurons (solid line) exceeds a threshold (dashed line). Avalanche duration is the number of time steps included in an avalanche, while avalanche size is the number of spikes included in an avalanche. **(b)** The probability distributions of avalanche duration and size under different I/E weight ratios. The color stands for different I/E weight ratios identical to the color in **c** and **d**. Vertical axis is logarithmic with the scale bar showing 3 orders of magnitude. Distributions are shifted vertically for comparison. **(c)** *κ*_1.7_ for avalanche duration as a function of I/E weight ratio and **(d)***κ*_1.5_ for avalanche size as a function of I/E weight ratio. *κ*_*ϵ*_ measures the deviation of a distribution from the power-law distribution with exponent *E*. For each I/E, we generated simulated realizations (dots) and averaged them to obtain the mean (solid line).

As discussed above, we constrain all of our models (all I/E ratios) such that the largest eigenvalue of the weight matrix is 1. This ensures that, on average, the population firing rate *S*(*t*) is stable, not growing or decaying over time, i.e. 〈*S*(*t* + 1)*/S*(*t*)〉_*t*_ = 1. Nonetheless, the temporal fluctuations in population firing rate can be quite large, particularly for low I/E. One way to take a closer look at why population firing fluctuates is to examine the ‘branching function’ developed in previous work [30]. The branching function Λ(*S*) is defined as the expected value of *S*(*t* + 1)*/S*(*t*) conditioned on the level of activity *S*. As shown in Fig 4a, for high I/E weight ratios, the branching functions cross Λ(*S*) = 1 at a certain point (red and yellow lines), which means the system is only stable at a certain level of activity and fluctuates closely around it. In contrast, for low I/E weight ratios, the branching functions have a wide range near Λ(*S*) = 1, which means the system is able to wander through various population firing rates. We define the ‘critical range’ *ρ* to measure the range of the branching function that stays within a certain distance of 1. Fig 4b shows that the critical range *ρ* is high when the I/E weight ratio is low, and it sharply decreases when the I/E weight ratio exceeds 2. Thus, the large fluctuations in population firing rate for low I/E are due to the large critical range, while small fluctuations at high I/E are due to a small critical range. This further clarifies how the network dynamics depend on I/E.

**Fig 4.**
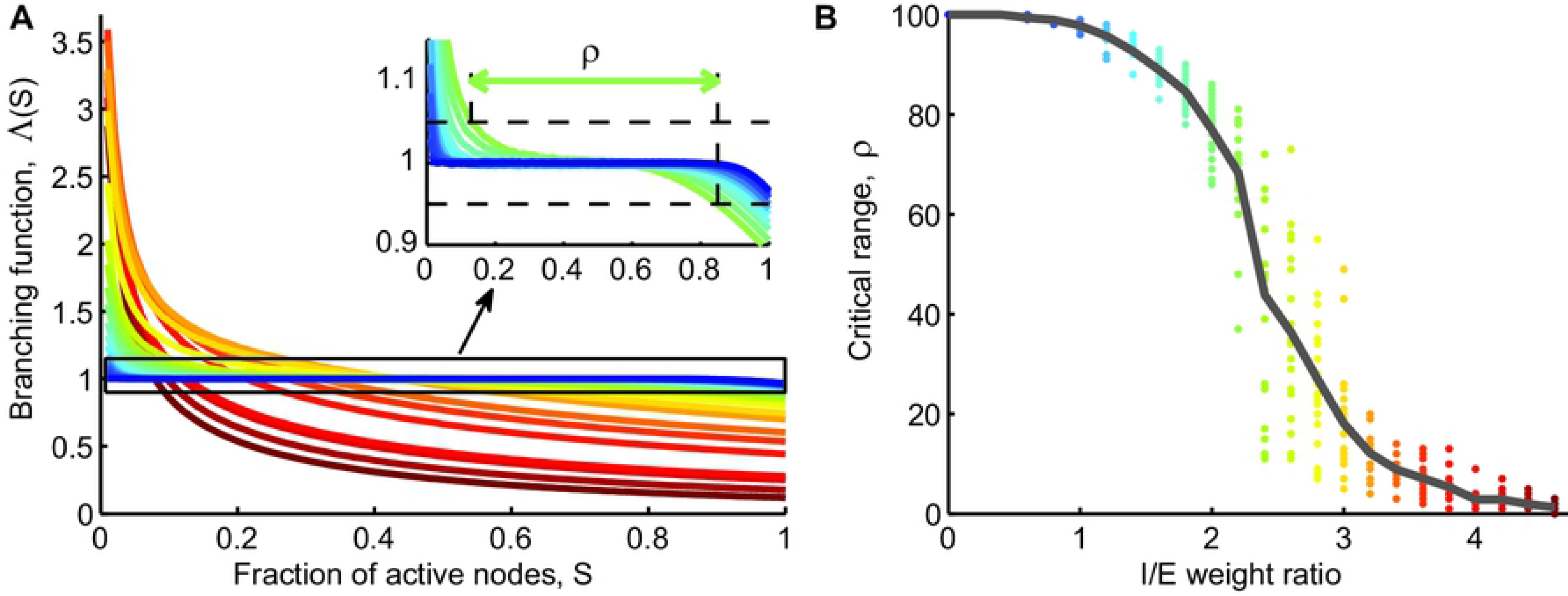
The critical range of branching function is wide under low I/E weight ratios and is narrow under high I/E weight ratios. **(a)** Branching functions under different I/E weight ratios. The color stands for different I/E weight ratios identical to the color in **b**. Inset: branching functions when the I/E weight ratio is less than 2.4. The critical range *ρ* is defined as the range between Λ = 1.05 and Λ = 0.95. **(b)** Critical range as a function of the I/E weight ratio. For each I/E, we generated simulated realizations (dots) and averaged them to obtain the mean (solid line).

Considering the avalanche distributions and branching functions, we can confidently conclude that our model operates near criticality for low I/E. Next, we sought additional evidence to support the apparent possibility that high I/E corresponds with the chaotic balanced state. Perhaps the most essential property of the chaotic balanced state is that neurons should fire asynchronously; neurons should be weakly correlated. Thus, we next examined the population-averaged input cross-correlogram (CCG) for different I/E weight ratios (Fig 5a). When the I/E weight ratio is low, CCGs of excitatory, inhibitory and total inputs are all high. As the I/E weight ratio increases, CCGs decrease. This decrease is most prominent at non-zero delays and for the total input CCG. When the I/E weight ratio is high, although the excitatory and inhibitory input CCGs remain relatively high at zero delay, the total input CCG is weak due to tight balance between excitatory and inhibitory inputs. We quantify asynchrony based on decreases in temporal and cross-neuron correlations. For this, we define *η* to be inversely proportional to the area under CCG for total input (normalized as defined the Methods). As shown in Fig 5b, asynchrony *η* sharply increases when the I/E weight ratio goes beyond 2, and reaches a high value when the I/E weight ratios is near 4. Therefore, strong inhibition makes excitatory and inhibitory inputs balanced and leads to an asynchronous activity. This is in accordance with the result in previous work that the population-averaged firing correlation is weak when inhibition is strong and fast [17].

**Fig 5.**
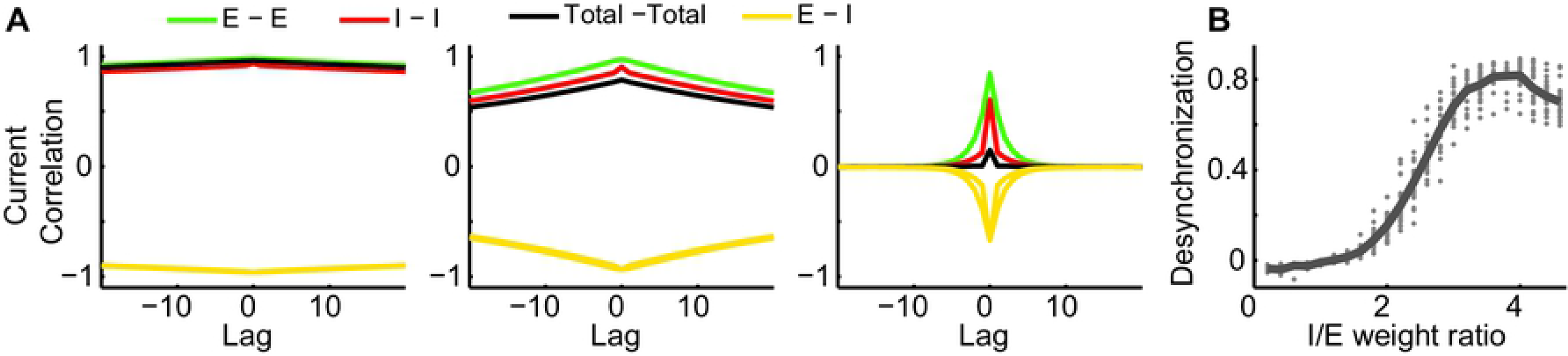
Asynchrony increases as the I/E weight ratio increases. **(a)** Population-averaged cross corelograms (CCGs) of total synaptic inputs (black), excitatory inputs (green) and inhibitory inputs (red) when the I/E weight ratios are 1 (left), 2 (middle) and 4 (right). Yellow reveals correlations between excitation and inhibition. **(b)** *η* as a function of I/E weight ratio. *η* measures the level of desychronization. For each I/E, we generated simulated realizations (dots) and averaged them to obtain the mean (solid line).

Another important feature found in chaotic balanched networks is a particular scaling relationship between the strength of synaptic weights *J* and the number of connections per neuron *K*. Specifically, *J* scales inversely with the square root of *K*, 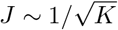 [21, 32]. We next examined if this scaling rule emerges as we increased I/E from low to high (see Methods). We found that for different I/E weight ratios, the synaptic strength *J* as a function of *K* are power-law distributions with different scaling exponents (Fig 6a). By curve fitting, we found a continuum of scaling exponents from low to high I/E weight ratios (Fig 6b). As shown in Fig 6c, when the I/E weight ratio is low, the exponent is near *−*1. When the I/E weight ratio is high, the exponent increases to around *−*0.5. Thus, we conclude that at high I/E, our model confirms the 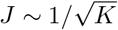 scaling expected for chaotic balanced networks. Considering together the tight balance of E and I inputs (Fig 2), the asynchronous firing (Fig 5), and the synaptic scaling (Fig 6), we conclude that high I/E in our model is consistent with the chaotic balanced regime.

**Fig 6.**
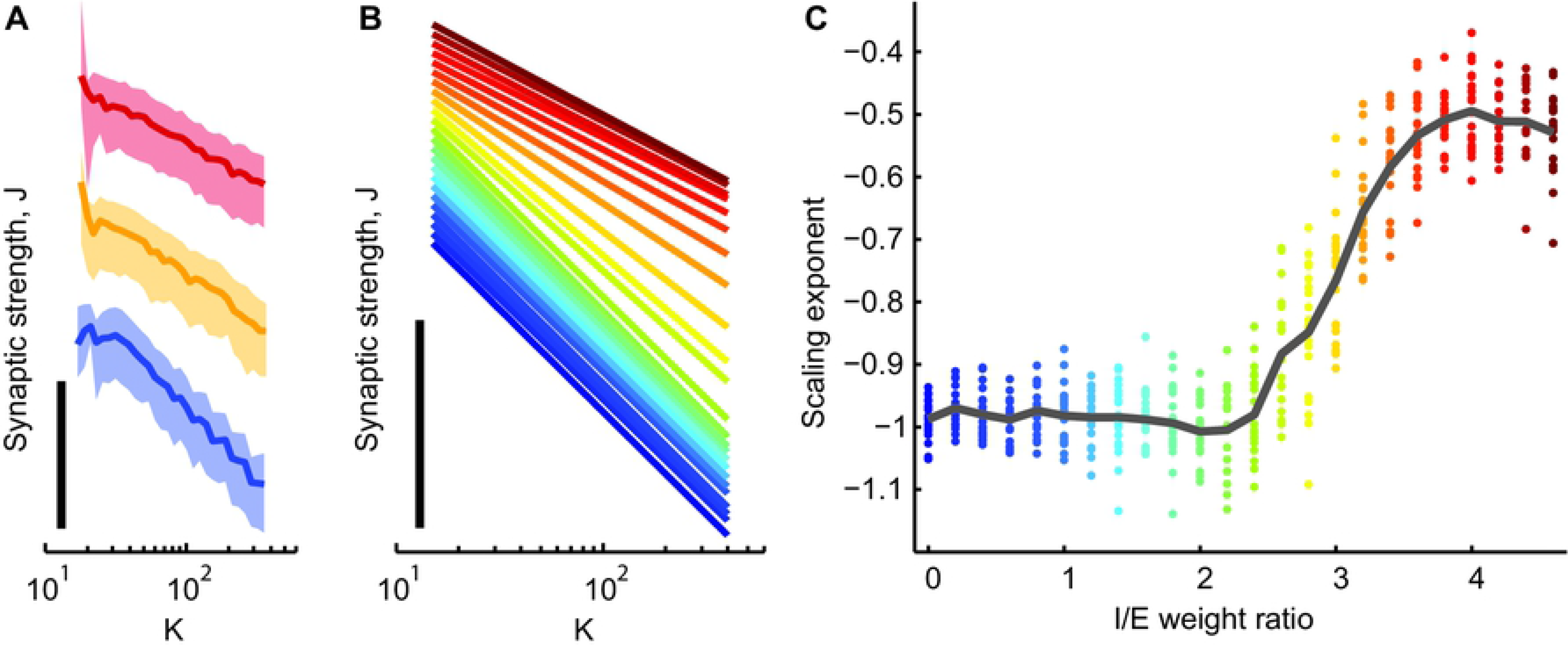
Synaptic strength scales with the number of connections as 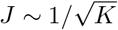 when the I/E weight ratio is high. **(a)** Synaptic strength *J* vs. the number of connections *K* when the I/E weight ratios are 0.6 (blue), 3.2 (orange) and 4.2 (red). The solid line shows the median and the shade shows the upper and lower quartiles. **(b)** *J* as a function of *K* using the best-fitted scaling exponents under different I/E weight ratios. The color stands for different I/E weight ratios identical to the color in **c**. **(c)** Best-fitted scaling exponents as a function of I/E weight ratio. For each I/E, we generated simulated realizations (dots) and averaged them to obtain the mean (solid line).

## Discussion

We have shown that the population activity of neural networks can vary dramatically depending how excitation and inhibition are balanced. If weak excitation is balanced by weak inhibition, we found that the dynamics exhibit large fluctuations and rather coordinated activity. If stronger inhibition balances stronger excitation in a higher “tension” balance, we found that the dynamics are asynchronous and steady, consistent with the chaotic balanced state. Our work establishes a simple bridge between two previously discrepant views of cortical population dynamics. Our findings suggest that the same network could be tuned from criticality to the chaotic balanced, by simply strengthening inhibition and excitation.

One interesting hypothesis that emerges from our work concerns metabolic efficiency. First, we note that maintaining a “strong” synapse depends on metabolically expensive biophysical mechanisms - greater presynaptic vesicle pool, greater density of postsynaptic receptors, etc. Since the high I/E regime and the low I/E regime have similar firing rates, it stands to reason that the strong synapses of the high I/E scenario would consume more metabolic resources than the lower I/E scenario. Moreover, the critical dynamics we observed at low I/E are associated with a number of functional benefits [10]. On the other hand, the lower fluctuations found in the high I/E regime may be beneficial for functions that require lower “noise” [21, 23]. When low noise is not required, perhaps the brain could tune itself to the low I/E regime where energy consumption is less. This is consistent with the observation that resting, awake animals tend to exhibit greater fluctuations in population activity compared to alert, active animals. Given the high firing rates and high noise of intermediate I/E (around 2), this regime may be less useful and more energetically costly. Experimental tests of this idea would be challenging, requiring comparisons of synaptic strengths across behavioral states. We would predict that alert, active states would exhibit stronger synapses (i.e. excitatory and ihibitory postsynaptic potentials that are larger in magnitude) than those found in quiescent, resting states.

Our findings suggest that a single cortical network can shift between two dramatically different dynamical regimes that have traditionally been viewed as incompatible: criticality and the chaotic balanced state. By bridging the gap between these two points of view, we are optimistic that our results help resolve the debate over what kinds of dynamical regimes can manifest in the cortex.

## Methods

### Network

We construct the recurrent network with *N* = 1000 integrate-and-fire neurons, where a fraction *α* = 0.2 of them are inhibitory. Each neuron randomly links to other neurons with a probability *p* = 0.2. The strength of excitatory links is uniformly distributed in [0*, J*_E_], while the strength of inhibitory links is uniformly distributed in [0*, J*_I_]. Thus, the synaptic connections of the network can be presented by a weighted matrix *J*, where the element *J*_*ij*_ stands for the strength of connection from neuron *j* to neuron *i*. *J*_*ij*_ is positive for an excitatory connection and negative for an inhibitory connection, while *J*_*ij*_ is 0 when there is no connection. We then normalize the matrix with the absolute value of its largest eigenvalue, so that the value of *J*_I_ and *J*_E_ does not affect the properties of the network independently, whereas the ratio of *J*_I_ and *J*_E_ matters. We define the variable *J*_I_/*J*_E_ as the I/E weight ratio, and let it vary in [0, 4.6] to represent different levels of inhibition.

### Dynamics

We apply probabilistic integrate-and-fire dynamics on the recurrent network. The state of each neuron is binary, either 1 or 0 corresponding to active or quiescent, respectively. At each time step *t*, the probability of neuron *i* being active depends on two independent factors: a probability *p*_*i*_ due to synaptic inputs from other neurons within the network and a probability *p*_ext_ due to external inputs or spontaneous firing.

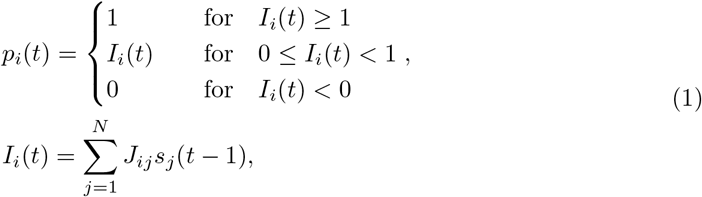

where *s*_*j*_ (*t* − 1) is the state of neuron *j* at time step *t −* 1, and *I*_*i*_(*t*) represents the total synaptic input to neuron *i* from other neurons at time step *t*. *p*_ext_ is set as 0.005*/N*, which corresponds to 1 externally-driven spike every 200 time steps over the whole network on average. We run the dynamics on the networks in Matlab 2010a.

### Synaptic input

The total synaptic input *I*_*i*_(*t*) can be separately examined in two parts: excitatory synaptic input 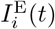 where only positive connections *J*_*ij*_ > 0 count, and inhibitory synaptic input 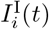 where only negative connections *J*_*ij*_ < 0 count. We define the E/I tension *T* to measure how tightly the 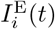 and 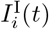 are balanced. The E/I tension of neuron *i* is defined as

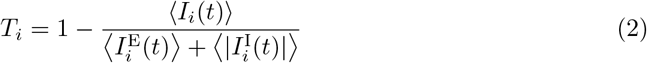

where 〈·〉 indicates time average. Then, the E/I tension *T* of the recurrent network is the average of *T*_*i*_ over all neurons.

### Avalanche distribution

The threshold for avalanches is determined at the level *S*\* when the branching function Λ(*S*\*) = 1.05, which is used in previous works [30]. We use *κ*_*ϵ*_ to measure how much the avalanche duration and size distributions deviate from power-law distributions [14]. *κ*_*ϵ*_ is defined as

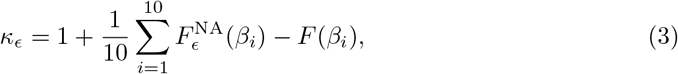

where 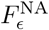 is the cumulative distribution function (CDF) of the reference power-law distribution with exponent *ϵ*, and *F* is the CDF of the avalanche duration or size. From the definition, *κ*_*ϵ*_ is close to 1 if the measured distribution well matches the reference power-law distribution, while either greater or smaller value of *κ*_*ϵ*_ means deviation from the reference power-law distribution. We use *ϵ* = 1.7 for avalanche duration and *ϵ* = 1.5 for avalanche size, which are the best-fitted exponents when there is no inhibitory neurons. We take *β*_*i*_ as a representative sample of 10 logarithmically spaced points along the measured distribution.

### Branching function

Branching function Λ(*S*) is defined as the expected value of *S*(*t* + 1)*/S*(*t*) under a certain level of activity *S*:

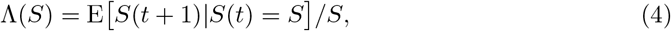

where *S*(*t*) is the number of active neurons in the time step *t*. Since not all values of S occur naturally in the models, we obtain a complete Λ(*S*) numerically, by simulating two time steps with the model many times for each possible S. Λ(*S*) *>* 1 means the system tends to have a higher level of activity, while Λ(*S*) < 1 means the system tends to have a lower level of activity. We use the critical range *ρ* = *S*_2_ *− S*_1_ to measure how long the branching function stays near 1, where *S*_1_ and *S*_2_ are the levels of activities when Λ(*S*_1_) = 1.05 and Λ(*S*_2_) = 0.95, as shown in Fig 3a insert.

### Input cross-correlogram (CCG)

We plot the input CCGs by calculating the population-averaged cross correlations of synaptic inputs with time lags from *−*20 to 20. To measure the desynchronization of the total synaptic input from the synchronized excitatory and inhibitory synaptic inputs, we define *η* as

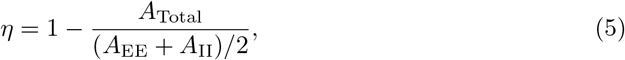

where *A*_Total_ is the area under total synaptic input CCG, and *A*_EE_ and *A*_II_ are the area under excitatory and inhibitory synaptic input CCGs, respectively.

### *J-K* scaling

To see the *J-K* scaling relationship, we vary the network size *N* logarithmically from 200 to 2000 while keeping all the other settings in our model the same. This generates a set of networks with different network sizes under a certain I/E weight ratio. Since the probability of connections *p* = 0.2 remains the same, the neurons in different networks have different number of connections as 〈*K*〉 = *Np*. Then, we take 200 neurons from each network to ensure the statistic is not biased. We fit selected samples to the function 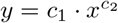, where *y* is the synaptic strength *J* and *x* is the number of the connections *K*, and find the best-fitted parameter *c*_2_ as the scaling exponent.

